# M3NetFlow: A novel multi-scale multi-hop graph AI model for integrative multi-omic data analysis

**DOI:** 10.1101/2023.06.15.545130

**Authors:** Heming Zhang, S. Peter Goedegebuure, Li Ding, David DeNardo, Ryan C. Fields, Yixin Chen, Philip Payne, Fuhai Li

## Abstract

**Summary:** Multi-omic data-driven studies, characterizing complex disease signaling system from multiple levels, are at the forefront of precision medicine and healthcare. The integration and interpretation of multi-omic data are essential for identifying molecular targets and deciphering core signaling pathways of complex diseases. However, it remains an open problem due the large number of biomarkers and complex interactions among them. In this study, we propose a novel Multi-scale Multi-hop Multi-omic graph model, *M3NetFlow*, to facilitate generic multi-omic data analysis to rank targets and infer core signaling flows/pathways. To evaluate M3NetFlow, we applied it in two independent multi-omic case studies: 1) uncovering mechanisms of synergistic drug combination response (defined as anchor-target guided learning), and 2) identifying biomarkers and pathways of Alzheimer ‘s disease (AD). The evaluation and comparison results showed *M3NetFlow* achieves the best prediction accuracy (accurate), and identifies a set of essential targets and core signaling pathways (interpretable). The model can be directly applied to other multi-omic data-driven studies. The code is publicly accessible at: https://github.com/FuhaiLiAiLab/M3NetFlow

## 1. Introduction

Multi-omic data-driven studies are at the forefront of precision medicine and healthcare. Recently, multi-omic datasets, like genetic, epigenetic, transcriptomic, and proteomic, have been being generated to characterize dysfunctional biological processes and signaling pathways from multiple levels/views, and to elucidate the panoramic view of the disease pathogenesis ^1–5^. For example, The Cancer Genome Atlas (TCGA) program have generated multi-omic datasets of over 20,000 samples spanning 33 cancer types, to understand the key molecular targets and signaling pathways of cancer ^6^. Moreover, the multi-omic data of >10,000 cancer cell lines were profiled in the Cancer Cell Line Encyclopedia (CCLE) project, which are valuable to investigate the mechanism of cancer response to given drugs and drug combinations^7^. In addition, the multi-omic data of Alzheimer’s disease (AD) are generated and publicly available in the ROSMAP^8^ project to uncover the pathogenesis of AD. Also, the exceptional longevity (EL), like the Long-Life Family Study (LLFS) project, have been generating multi-omic data^9–11^ to identify protective biomarkers and pathways for long and healthy life. The multi-omic data are valuable and essential for understanding the key molecular targets and mechanisms of diseases, identifying novel therapeutic targets, predicting effect drugs and drug cocktails to guide the development of precision medicine.

However, it remains an open problem and a challenging task for integrative and interpretable omic data analysis, though a set of computational models has been proposed. A comprehensive review of existing multi-omic data integration analysis models was reported^12^. Specifically, these models were clustered into a few categories, like similarity, correlation, Bayesian, multivariate, fusion and network-based models. The PAthway Representation and Analysis by Direct Inference on Graphical Models (PARADIGM^13^) is one of the most widely used methods among these traditional computational methods. Aside from that, the NMTF-based method called iCell^14^ was also proposed to integrate the multi-omic data and only the direct connection was considered. The timeOmics^15^ was recently developed to identify signaling patterns over time of biomarkers in multi-omic data, allowing for the analysis of time-series data in biological systems.

### Problem formulation

The problem to be tackled in this study using graph AI models is to identify or rank disease or drug response associated molecular biomarkers from a large number of multi-omic features, and infer the potential signaling network among the selected biomarkers. The two specific biological problems to be tackled using the graph AI models are as follows. The first task is an anchor-biomarker (like known molecular targets, or drug targets) guided learning, e.g., to identify targets and pathways regulating drug cocktail response by integrating multi-omic data of cells and drug targets. The input data of the first case study is the multi-omic profiling of individual cell lines, kegg signaling pathways (graph), drug targets, and synergistic scores of a set of drug combinations. The output of the model is a set of top-ranked biomarkers and signaling pathways linking to the given anchor-biomarkers (information flowing to the anchor-biomarkers), which are informative to predict and explain the mechanism of drug combination response. In addition to drug-targets, the anchor-biomarkers can be generic given targets of interested to study the up- or down-stream signaling pathways of the anchor-biomarkers. The second task is to identify disease associated biomarkers and signaling pathways of Alzheimer’s disease (AD). It is a generic biomarker ranking and pathway inference problem without the guide/constraint of anchor-biomarkers. The input is multi-omic data, like the AD and control/normal samples and kegg signaling pathways (graph). The output of the model is a set of top-ranked biomarkers and signaling pathways, which are informative to classify AD from normal samples explaining the pathogenesis of AD. The two biological problems/case studies represent two of the most needed multi-omic data analysis tasks.

### Related work

Graph Neural Networks (GNNs) have gained prominence due to their capability to model relationships within graph-structured data^16–19^. And numerous studies have applied the Graph Neural Network (GNN) with the integration of the multi-omic data. MOGONET^20^ initially creates similarity graphs among samples by leveraging each omic data, then employs a Graph Convolutional Network (GCN^16^) to learn a label distribution from each omic data independently. Subsequently, a Cross-omic discovery tensor is implemented to refine the prediction by learning the dependency among multi-omic data. MoGCN^21^ adopts a similar approach by constructing a patient similarity network using multi-omic data and then using GCN to predict the cancer subtype of patients. GCN-SC^22^ utilizes a GCN to combine single-cell multi-omic data derived from varying sequencing methodologies. MOGCL^23^ takes this further by exploiting the potency of graph contrastive learning to pretrain the GCN on the multi-omic dataset, thereby achieving impressive results in downstream tasks with fine-tuning. Nevertheless, none of the aforementioned techniques contemplate incorporating structured signaling data like KEGG into the model. Moreover, general GNN models are limited by their expression power, i.e., the low-pass filtering or over-smoothing issues, which hampers their ability to incorporate many layers. The over-smoothing problem was firstly mentioned by extending the propagation layers in GCN^24^. Moreover, theoretical papers using Dirichlet energy showed diminished discriminative power by increasing the propagation layers^25^. And multiple attempts were made to compare the expressive power of the GCNs^19,26^, and it is shown that WL subtree kernel^27^ is insufficient for capturing the graph structure. Hence, to improve the expression powerful of GNN, the ***K***-hop information of local substructure was considered in various recent research^28–33^. However, none of these studies was specifically designed to well integrate the biological regulatory network and provide the interpretation with important edges and nodes.

In this study, we present a novel graph AI model, named ***M3NetFlow (Multi-scale, Multi-Hop, Multi-omic NETwork Flow inference)*** to address the challenges mentioned above. Specifically, using an attention mechanism, our approach tackles these challenges by first incorporating local ***K***-hop information within each subgraph by integrating the gene regulatory network such as KEGG^34^ and BioGrid^35,36^. Subsequently, we establish global-level bi-directional propagation to facilitate message passing between genes/proteins. In the interpretation phase, we leverage the attention mechanism to aggregate the weights of all connected paths for genes, and a reweighting process inspired by (Term Frequency – Inverse Document Frequency) TF-IDF^37^ is employed to redistribute the importance scores of genes across different cell lines. Afterward, a visualization tool, ***NetFlowVis***, was developed for the better analysis of targets and signaling pathways of drugs and drug combinations. To assess and demonstrate the effectiveness of our proposed model, ***M3NetFlow***, applied it in two independent multi-omic case studies: 1) uncovering mechanisms of synergistic drug combination response (defined as anchor-target guided learning), and 2) identifying biomarkers and pathways of Alzheimer’s disease (AD). The evaluation and comparison results showed *M3NetFlow* achieves the best Pearson correlation or prediction accuracy (accurate), and identifies a set of essential targets and core signaling pathways (interpretable, which can be directly applied to other multi-omic data-driven studies.

## 2 Methodology

**Table 1** provided the links from which four types of datasets, drug combination effects, multi-omic data, gene-gene interactions, and drug-gene interactions, were collected to predict drug combinations for cancer cell lines. What’s more, to investigate Alzheimer’s disease, we also obtained corresponding multi-omic datasets and clinical dataset from publicly available sources, particularly the ROSMAP datasets (see **Table 2**). Comprehensive details regarding the data and the preprocessing results are thoroughly documented in Appendix Section A. This section provides an in-depth explanation of the methodologies employed, as well as the specific parameters and outcomes obtained during the preprocessing stage.

**Table 1.**
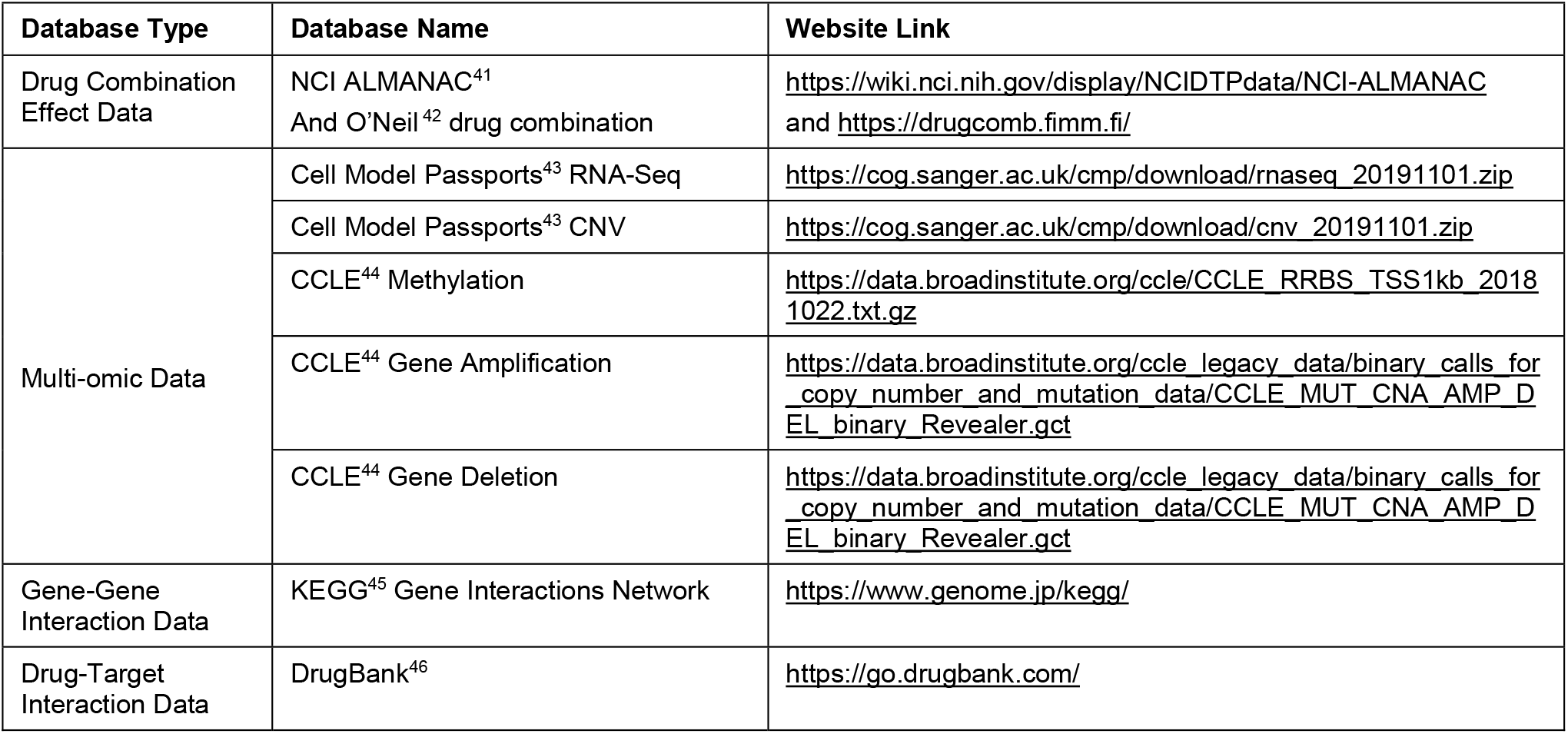
Multi-omic data and drug combination screening data of cancer cell lines.

**Table 2.** Multi-omic data and clinical data from ROSMAP database resources.

| Database Type | Database Name | Website Link |
| --- | --- | --- |
| Clinical Data | ROSMAP_clinical | https://www.synapse.org/#!Synapse:syn3191087 |
| Multi-omic Data | ROSMAP_arrayMethylation_imputed | https://www.synapse.org/#!Synapse:syn3168763 |
|  | ROSMAP.CNV.Matrix(Mutation) | https://www.synapse.org/#!Synapse:syn26263118 |
|  | ROSMAP_RNAseq_FPKM_gene | https://www.synapse.org/#!Synapse:syn3505720 |
|  | C2.median_polish_corrected_log2(Proteomic) | https://www.synapse.org/#!Synapse:syn21266454 |
| Mapping Data | GEO GPL16304 Platform | https://www.ncbi.nlm.nih.gov/geo/query/acc.cgi?acc=GPL16304 |

### 2.1 Model Architecture of M3NetFlow

Figure 1. shows the schematic architecture of the proposed ***M3NetFlow*** model. The model input parameters are 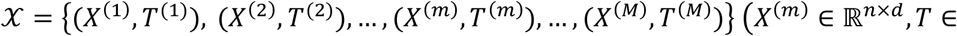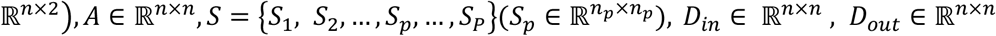, where *M* represents number of data points in the drug screening dataset. To predict the drug combination effect scores *Y* (*Y* ∈ ℝ^M×1^), we would like to build up the machine learning model *f*(·) with *f*(*X, A, S, D*_i_, *D*_out_) = *Y*, where *𝒳* denotes all of the data points in the dataset and (*X*^(*m*)^, *T*^(*m*)^) is *m*-th data points in the dataset, where *X*^(*m*)^ denotes the node features matrix with *n* nodes of *d* features and *T*^(*m*)^ denotes the one-hot encoding of the number of drugs targeted on those *n* nodes. The matrix *A* is the adjacency matrix that demonstrates the node-node interactions, and the element in adjacency matrix *A* such as *a*_*ij*_ indicates an edge from *i* to *j*, and *S* is a set of subgraphs that partition the whole graph adjacent matrix *A* into multiple subgraphs with 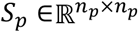 of nodes interactions between its internal *n*_*p*_ nodes and each subgraph has its own corresponding subgraph node feature matrix 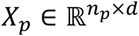. *D*_*in*_ is an in-degree diagonal matrix for nodes in directed graph, and *D*_*out*_ is an out-degree diagonal matrix for nodes in directed graph.

**Figure 1.**
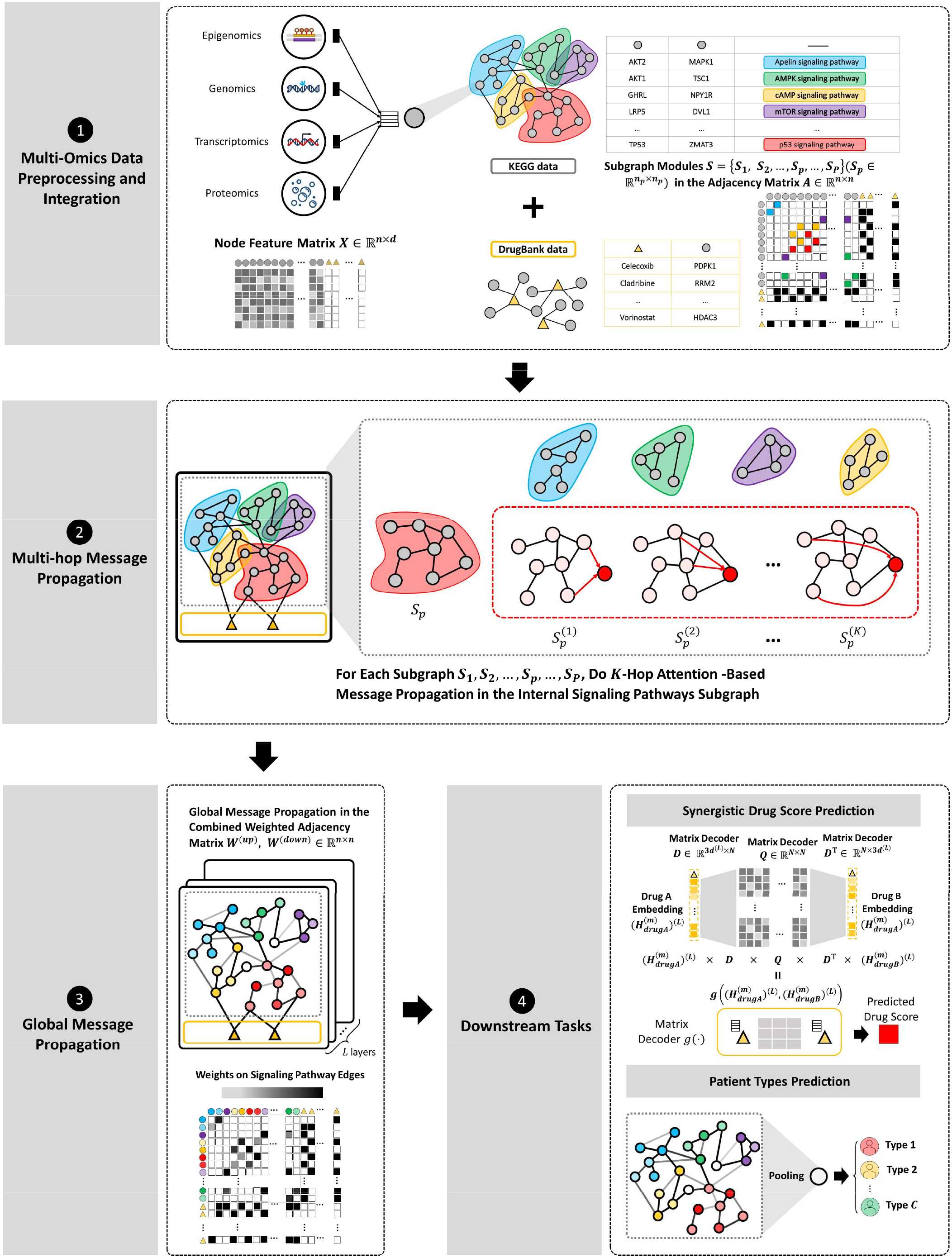
Model architecture of *M3NetFlow*. ➊Integrate the one-hot encoded and multi-omic features into a vector for each node. Afterwards, Merge drug-gene and gene-gene interactions into an adjacency matrix. ➋ Multi-hop attention-based propagation was performed in the subgraphs. ➌ Use combined weighted adjacency matrix for global signaling propagation.➍ Downstream tasks. (1) Decode the pair of drug nodes to predict the drug synergy scores. (2) Use pooling strategy to predict the patient outcomes.

#### Network Modular and Multi-hop Message Propagation

In the graph message passing stages of our architecture (see **Figure 1** step3 and step4), the multi-scale design, i.e., the local network module/subgraph module message passing stage for each signaling pathway, and the global message passing stage were designed. The multi-scale design will ensure the message fully interacts in the internal subgraph. Moreover, we opted for a *K*-hop attention-based graph neural network because it further allows for the consideration of longer distance information, which can be interpreted as signaling flow in a single message propagation layer. With those initial embedding features *X*^(*m*)^ ∈ ℝ^*n*×*d*^, (*m* = 1, 2, …, *M*), the *K* -hop attention-based graph neural network was built to incorporate the long-distance information for each subgraph with

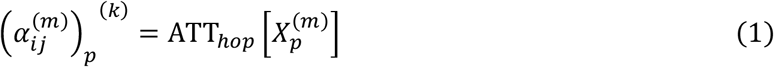

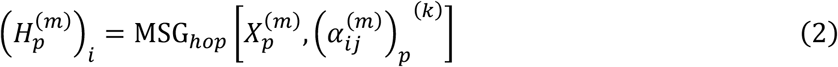

where 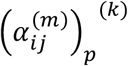 is the attention score between node *i* and node *j* in the *k* -th kop of the subgraph *S*_*p*_ for the data point *m* and the updated node feature 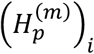 for node *i* will be generated via *K*-hop message propagation (see **Appendix B.1** for details).

#### Global Bi-directional Message Propagation

Following the message propagation in the multiple internal subgraphs, the global weighted bi-directional message propagation will be performed, where nodes-flow contains both ‘upstream-to-downstream’ (from up-stream signaling to drug targets) and ‘downstream-to-upstream’ (from drug targets to down-stream signaling) (see **Figure 1** step 3). Before the global level message propagation, the node feature for data point *m* in each subgraph 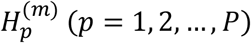 will be combined into a new unified node features matrix 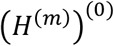 as the initial node features with

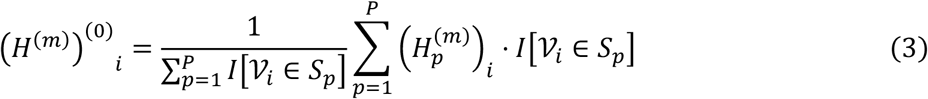

where *V*_*i*_ represents the node/vertex *i* in the graph and *I*[*V*_*i*_ ∈ *S*_*p*_] is the indicator function, whose value will be one if 𝒱_*i*_ ∈ *S*_*p*_. In the end, the initial node features for global level propagation will be 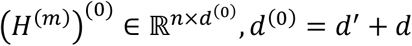, *d*^(0)^ = *d*^’^ + *d* . Then 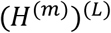 will be generated by weighted bi-directional message propagation via

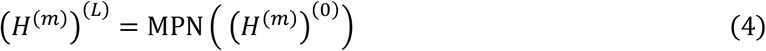

where MPN is the global bi-directional message propagation network (see **Appendix B.2** for details).

### 2.2 Downstream tasks

#### Drug combination response predictions by decoding drug node embeddings

After obtaining the embedded node features 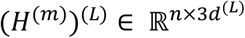 from the global message passing network, the features for drug A are represented as 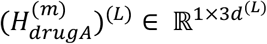 and the features for drug B are represented as 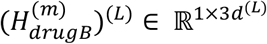 .Utilizing the decagon decoder^47^, the prediction of combo score will be calculated in the following equation:

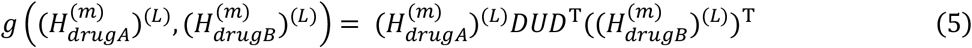

where 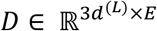 and *U* ∈ ℝ^*E*×*E*^ are trainable decoder matrices, as illustrated in Figure 1 from step 4.

#### Patient outcome predictions

With the embedded node features, the global mean pooling strategy was applied to predict the patient outcome with

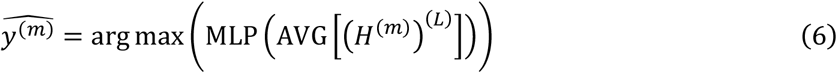

where 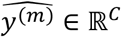 and *C* is the number of sample types (see Figure 1 step 4).

### 2.3 Interpretation using subgraph attention-based score

As aforementioned, we added the *K*-hop subgraph message propagation with attention in each subgraph, and the attentions or weights on each edge can potentially indicate and interpret the signaling flow on the signaling network to affect the drug combination response. Specifically, putting the attentions/weights on each edge will use the trained parameters *a* ∈ ℝ^2*d*’^and *W* ∈ ℝ^*d*×*d*’^. And for cell lines set, the set *𝒞* = {*C*_1_, *C*_2_, …, *C*_*r*_, …, *C*_*R*_} contains *R* = 39 cell lines; for ROSMAP AD dataset, *R* = 128, and we can choose the input node feature matrix within a specific cell line or sample to form the cell-line / sample specific attention matrix in *K*-hop by

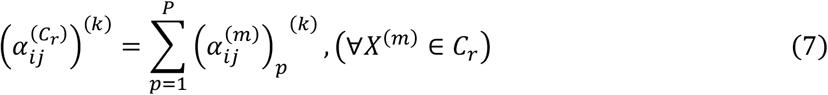

where 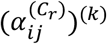 will be the element in 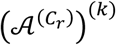, the *k*-hop attention matrix for cell line *C*_*r*_. And above formula shows the calculation for the 1-head attention matrix. For the multi-head attention mechanism, the averaged value will be used for the attention matrix.

## 3. Results

### 3.1 Experimental Setup

To validate the proposed model on drug prediction task, we implemented a 5-fold cross-validation approach. This method involved the use of a four-element vector, <*D*_*A*_, *D*_*B*_, *C*_*C*_, *S*_*ABC*_>, as the model input. In this tuple, *D*_*A*_ and *D*_*B*_ denote the specific drug combinations being analyzed. *C*_*C*_ represents the cell line name, which is integrated with multi-omic data to provide comprehensive contextual information. Lastly, *S*_*ABC*_ reflects the synergistic drug effect observed on the cell line *C*_*C*_. In total, there are 2788 model input data points for the NCI ALMANAC dataset and 1008 model input data points for the O’Neil dataset. In each fold model, 4 folds of the data points will be used as the training dataset, and the rest 1-fold will be used as the testing dataset. For those 1489 genes, their 8 features indicate the RNA sequence number, copy number variation, gene amplification, gene deletion, gene methylation maximum and minimum value and whether they have a connection spanning *D*_*A*_ and *D*_*B*_ respectively. Regarding the AD samples outcome prediction on ROSMAP dataset, we utilized 138 samples from the ROSMAP dataset, categorized by disease status (74 AD, 64 non-AD). To address the data imbalance, we performed downsampling for the classification task. For the AD vs. non-AD classification, we downsampled the AD samples to match the 64 non-AD samples, resulting in a dataset with 64 AD and 64 non-AD samples. Afterwards, 5-fold cross-validation was leveraged to evaluate the performance of our model. For those 2099 genes, their 10 features indicate the methylation values on upstream, distal promoters, proximal promoters, core promoters and downstream, the genetic mutations with (number of duplicates, deletions and mCNV), the gene expression values and protein expression values. For each drug pair, their 8 features in drug prediction task or 10 features in ROSMAP sample outcome prediction task were initiated with zeros. Furthermore, to indicate connections between nodes, adjacency matrices were also created. Those matrices were formed from the KEGG, which contains gene pairs with sources and destinations. For drug-gene edges, the connections are bidirectional, which means that in adjacency matrices, those elements are symmetric.

### 3.2 Hyperparameters

Subsequently, a model was developed by using pytorch and torch geometric. For both drug prediction and ROSMAP sample outcome prediction tasks, the learning rate started at 0.002 and was reduced equally within each batch for a certain epoch stage. And the epochs after 60 will keep the learning rate at 0.0001. Adam optimizer was chosen for optimization with eps=1e-7 and weight_decay=1e-20. We empirically set the ***K-hop Subgraph Message Propagation*** part with *K* = 3 and the ***Global Bi-directional Message Propagation*** part with *L* = 3 . Afterward, the feature dimensions will vary at the different layers and will be denoted by: (*d*^(1)^, *d*^(2)^, …, *d*^(*l*)^, …, *d*^(*L*)^). At the global message propagation part, the layer will concatenate biased node features and transformed node features of the previous layer for both upstream and downstream, generating concatenated dimensions for the output dims being 3 × *d*^(*l*)^ in *l*-th layer. Output dims of the previous layer served as the input dims for the current layer, as follows: (1) First layer (input dims, output dims): (*d*^(0)^, 3*d*^(1)^); (2) Second layer (input dims, output dims): (3*d*^(1)^, 3*d*^(2)^); (3) Third layer (input dims, output dims): (3*d*^(2)^, 3*d*^(3)^). The final embedded drug node dims were 3*d*^(3)^, (*L* = 3). For drug prediction task, the decoder trainable transformation matrix dims: 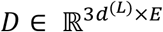 and *U* ∈ ℝ^*E*×*E*^ were used as trainable decoder matrices, with changeable parameters of *E* to adapt model performance and the model used *E* = 150 . In ROSMAP sample prediction task, the trainable graph mean pooling was leveraged to predict the sample outcome. As for the LeakyReLU function, the parameter α was set as 0.1 for both tasks.

### 3.3 *M3NetFlow* improves drug combination synergy and AD prediction accuracy

To evaluate the model performance in terms of synergy score prediction for drug combinations and predictions on ROSMAP AD samples, we conducted 5-fold cross-validation. As shown in **Table 3**, the average prediction (using the Pearson correlation coefficient), was about 61% Pearson correlation using the test data in the NCI ALMANAC dataset and was about 64% Pearson correlation using the test data in the O’Neil dataset. Regarding the ROSMAP dataset, the average prediction accuracy was about 66% using the test data in the ROSMAP dataset. These prediction results are comparable with existing deep learning models^38,39^. Moreover, we also compared our proposed model M3NetFlow with other deep learning models, which included the GCN^40^, Graph Attention network^41^ (GAT), UniMP^42^, MixHop^30^, Principal Neighborhood Aggregation^43^ (PNA) and GIN^44^. By checking the p values over 5-fold cross validation, the performances of the M3NetFlow have significant improvement over most of the GNN-based methods (see **Table 3** and **Figure 2a**).

**Table 3.**
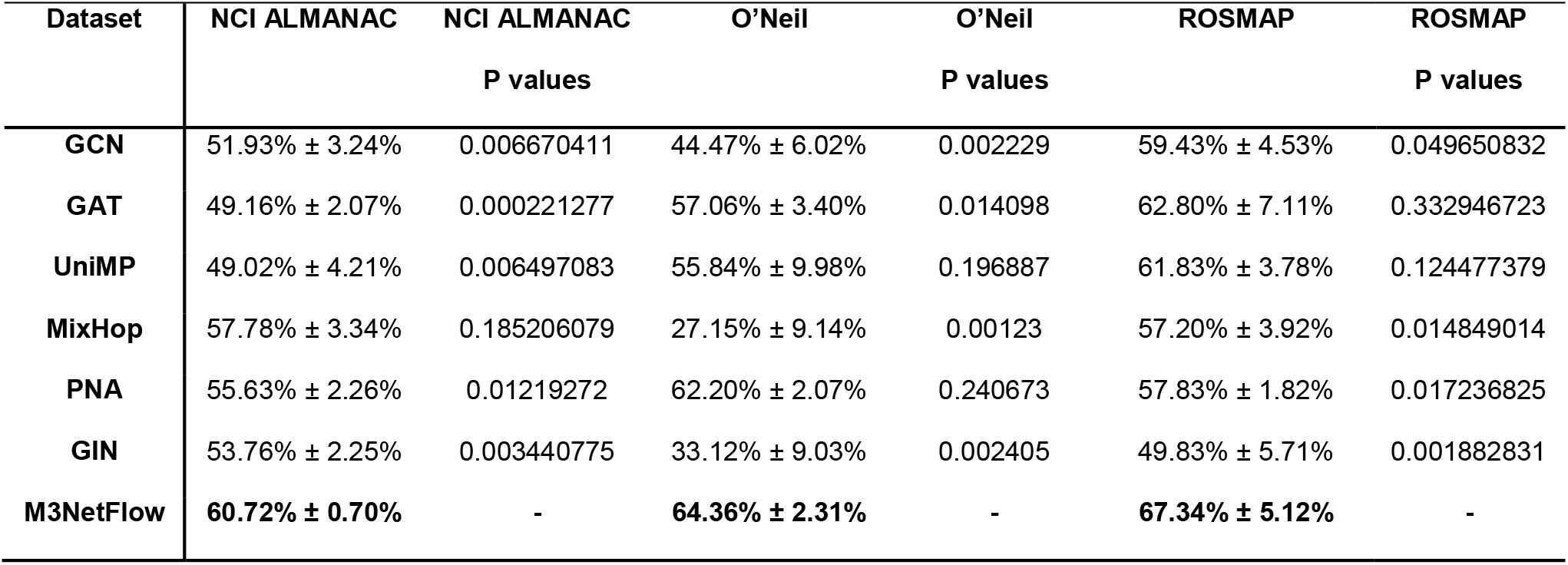
Model comparisons using average Pearson correlation and prediction accuracy of 5-fold cross-validation using NCI ALMANAC, O’Neil datasets and ROSMAP datasets.

**Figure 2.**
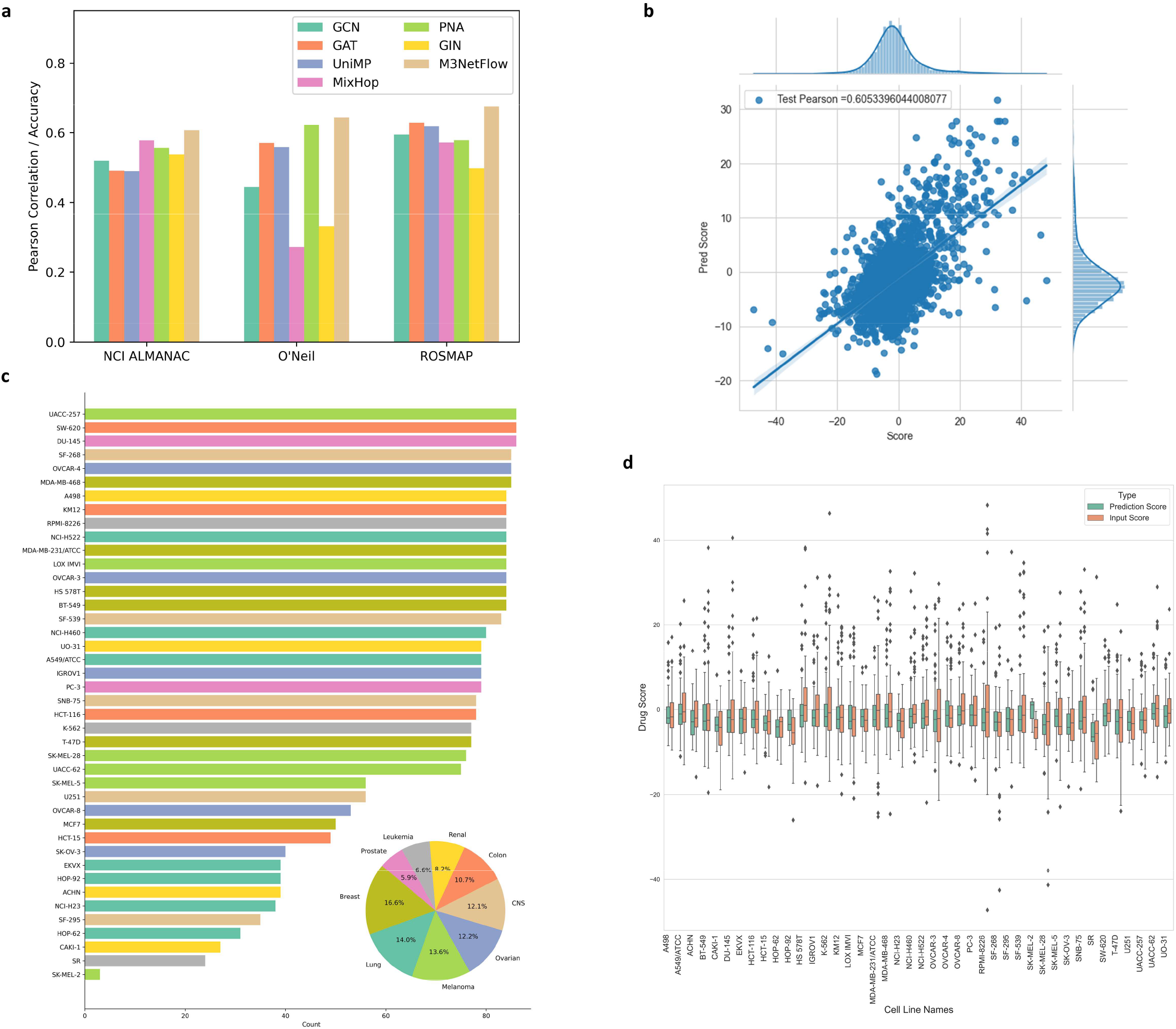
Model performance and overview of input dataset NCI ALMANAC, O’Neil (drug combination multi-omic data) and ROSMAP (AD multi-omic data). **a**. Pearson correlation comparisons for GCN, GAT, UniMP, MixHop, PNA, GIN and ***M3NetFlow*** models for NCI ALMANAC, O’Neil and ROSMAP dataset. **b**. Scatter plot of the model with data points in whole NCI ALMANAC dataset. **c**. Distributions of all cell lines in whole NCI ALMANAC dataset. **d**. Box plots across all cell lines in the whole NCI ALMANAC dataset.

### 3.4 *M3NetFlow* ranks important targets via attention score

Based on the attention matrix of each sample (like a cell line or an AD patient sample) between neighboring nodes (genes) on the graph in each fold of cross validation, the average attention matrix in was calculated. Therefore, the weighted importance of each node (gene) of each sample will be calculated based on the attention with

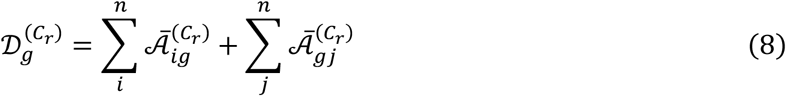

where 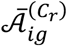 is the element of averaged 5-fold attention matrix in 1^st^ hop from a sample *C*_*R*_ in the *i*-th row and *j*-th column and 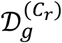 is the node importance score for gene *g* in the sample *C*_r r_Combining all the node importance vectors, the matrix *𝒟* ∈ ℝ^*n*×*R*^ will be generated, where n is the number of nodes, and R is the number of samples. However, some of genes may show the importance of relatively higher scores in each sample, which weakens the analysis of the sample-specific analysis. In this way, the idea of reweighting the gene importance score was created based on the TF-IDF to reweight the node importance in each sample with

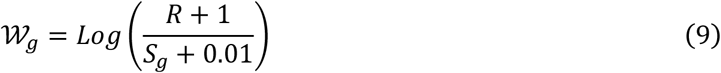

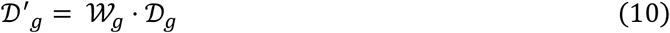

where *S*_g_ is the number of samples with a higher node importance score than the threshold *𝒮* in all samples. Here, the 95 percentile of node degree values in the whole matrix *𝒟* was used for *𝒮*. Finally, the reweighted node degree (importance score) matrix *𝒟*’ will be generated. Based on the reweighted matrix *𝒟*’, the node (gene) importance score will be calculated. The sum of importance score of individual genes across samples of disease will be used as the disease importance score of genes.

### 3.5 Case 1: Synergistic drug combinations are associated with the identified key targets

Based on the attention score calculated for each specific cell line in the section 3.4, the highest 5, and lowest 5 drug scores and their corresponding drugs and drug-targeted genes were collected for each cancer cell lines in NCI ALMANAC dataset (check **Figure 3a**). Comparing the targeted genes by drugs from the highest 5 drug scores and lowest drug scores, the differences are statistically significant with p values in more than half cell lines. Since the SK-MEL-2 cell line only has 3 drug screen data points and all of them are smaller than 0, this cell line was excluded from the statistical analysis. In total, there are 27 out of 41 (∼65%) cell lines have p values smaller than 0.1 (filled with deep orange color in **Table S1**) and 32 out of 41 (∼78%) cell lines have p value smaller than 0.3 (filled with light orange color in **Table S1**). And 37 out of 41 (∼90%) cell lines have higher gene importance scores on both mean and median values (filled with light green in **Table S1**). **Figures 4-6** provided more explicit evidence of a trend where genes targeted by drugs with higher scores exhibited higher node degrees (gene importance scores). In each cell line, the left boxplots compared the node degree distribution of genes targeted by the top 5 drugs with the highest scores against those targeted by the bottom 5 drugs with the lowest scores. The right boxplots in each cell line represented the overall comparisons between the top 5 and bottom 5 drug scores. Consequently, we could assign prior targets to genes with the highest node degree (gene importance scores) using our calculations. This allowed us to rank the genes in each cell line according to their node degree (importance scores), which can be found in **Table S2**. We generated a list of the top 20 genes in each cell line, available in **Table S3**. Subsequently, we identified overlapping genes among the cell lines of the same cancer type based on the top 20 genes in each line for cancer-specific analysis, as depicted in **Figure S1** from Appendix Section C.

**Figure 3.**
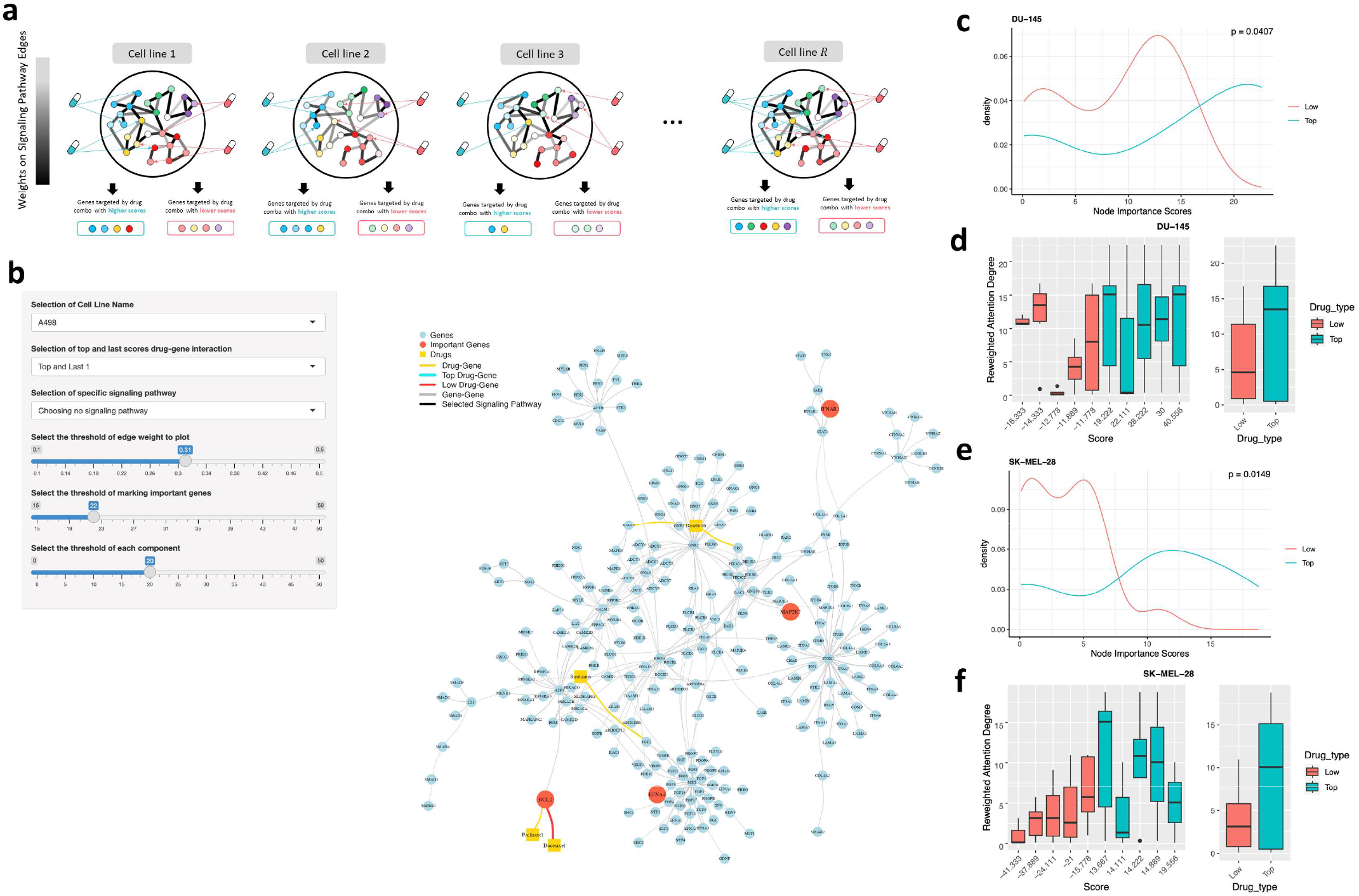
Validation and visualization through important node analysis **a**. Procedures of analyzing the important genes targeted by by the highest 5 and lowest 5 drug combinations **b**. Visualization tool ***NetFlowVis*** (check Appendix Section D for details of this tool) for core signaling network interactions of cell line DU-145 **c-f**. Density and box plots by comparing the genes targeted by the highest 5 and lowest 5 drug combinations in DU-145 and SK-MEL-28 cell lines.

**Figure 4.**
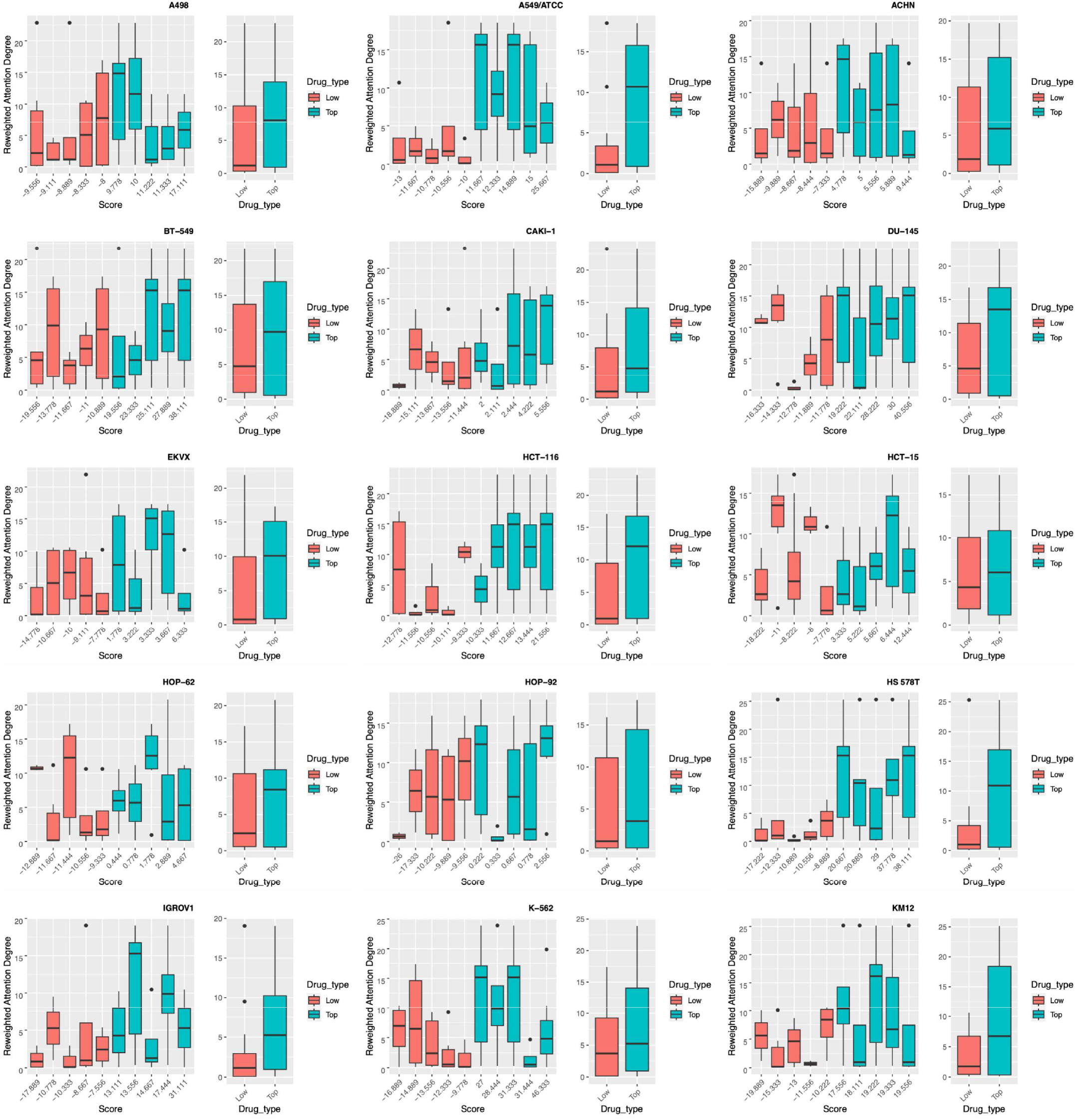
Cell line A498, A549/ATCC, ACHN, BT-549, CAKI-1, DU-145, EKVX, HCT-116, HCT-15, HOP-62, HOP-92, HS 578T, IGROV1, K-562, KM12 genes degree distribution targeted by highest 5 and lowest 5 drug scores.

**Figure 5.**
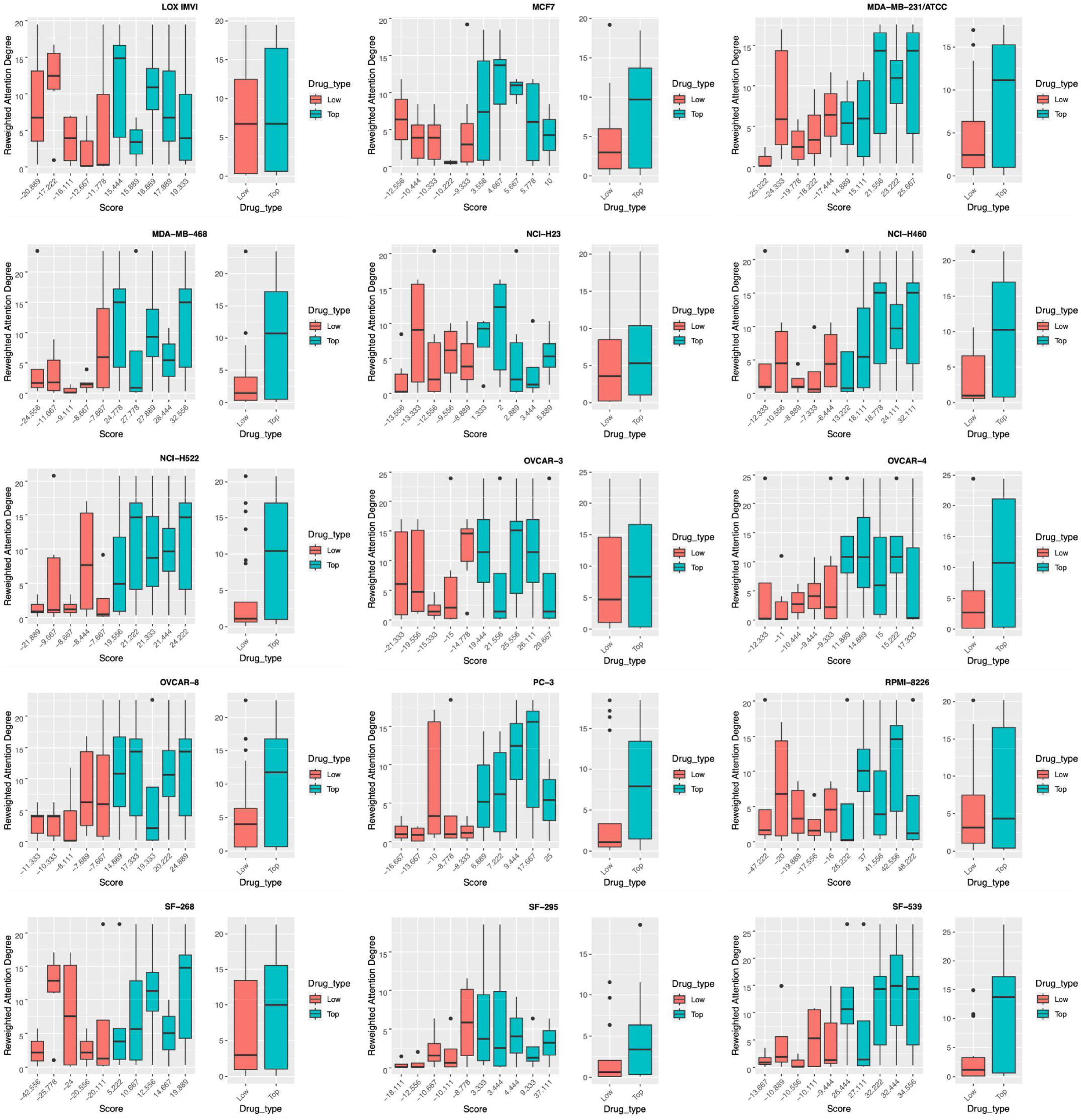
Cell line LOX IMVI, MCF7, MDA-MB-231/ATCC, MDA-MB-468, NCI-H23, HCI-H460, NCI-H522, OVCAR-3, OVCAR-4, OVCAR-8, PC-3, RPMI-8226, SF-268, SF-295, SF-539 genes degree distribution targeted by highest 5 and lowest 5 drug scores.

**Figure 6.**
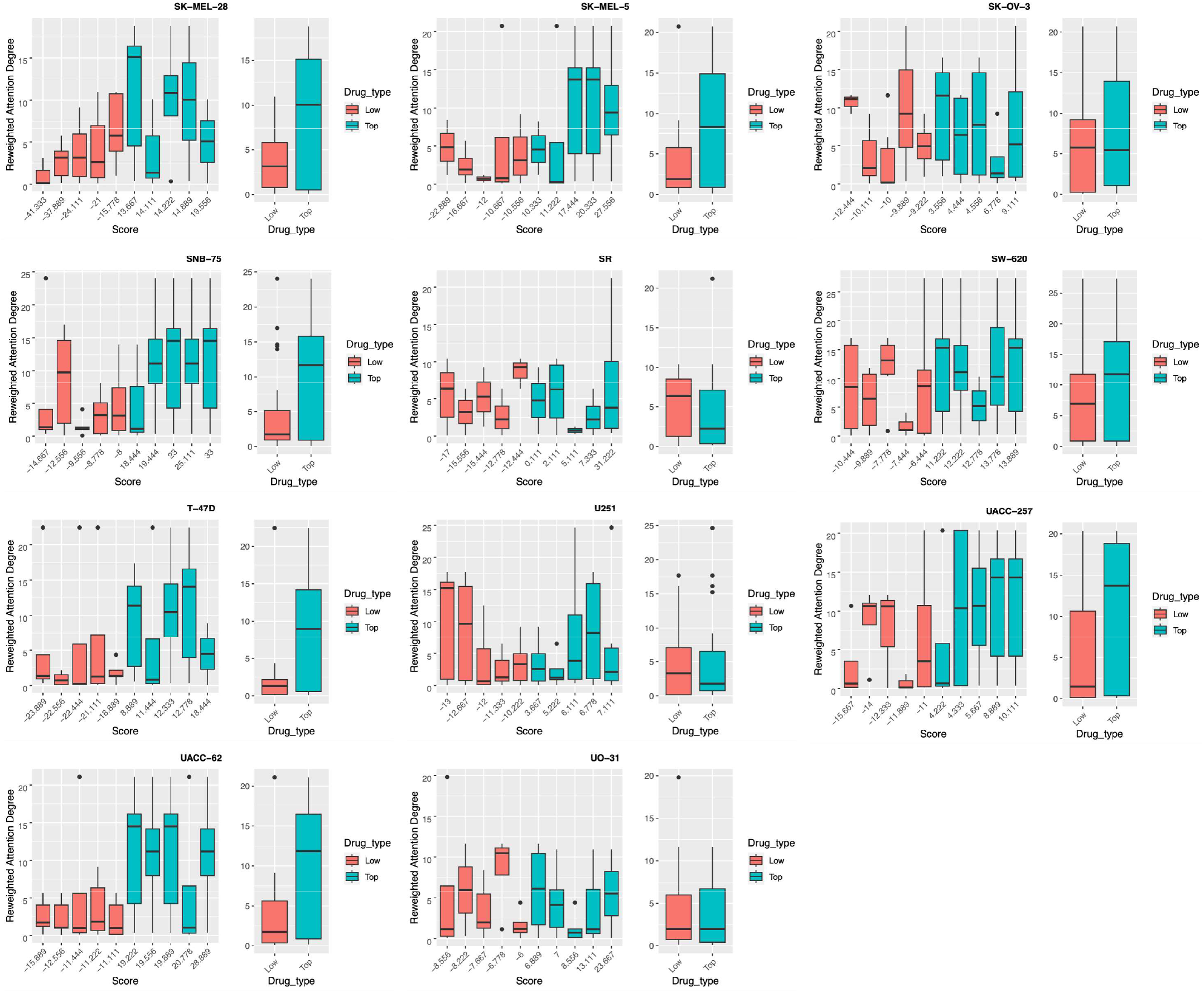
Cell line SK-MEL-28, SK-MEL-5, SK-OV-3, SNB-75, SR, SW-620, T-47D, U251, UACC-257, UACC-62, UO-31 genes degree distribution targeted by highest 5 and lowest 5 drug scores.

### 3.6 Case 2: Alzheimer’s disease associated biomarkers and pathways

Generating attention-based scores from ***M3NetFlow*** (see Section 3.4), the sample specific edge weight matrices will be aggregated by

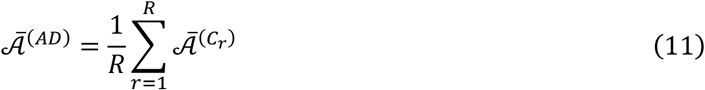

where *R* = 64 is the number of AD samples in the set *𝒞* and 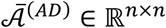 is the aggregated edge weight for AD patients. By setting the filters (edge threshold as 0.106 and a small component threshold as 15), we identified 100 potential important genes for AD. Among those genes, 28 genes are filtered out by setting the threshold of attention-based node weight as 2.0, and 15 of them with p values smaller than 0.1 in at least one of the 10 multi-omic features (see **Figure 7a-b**). To evaluate the top targets ranked by attention-based node weight, the top-ranked 28 AD associated biomarkers were further analyzed via pathway enrichment analysis. Interestingly, a set of AD associated signaling pathways are identified. As shown in **Figure 7c**, the top-ranked targets are involved in a set of signaling transduction pathways, which indicates the importance of these targets.

**Figure 7.**
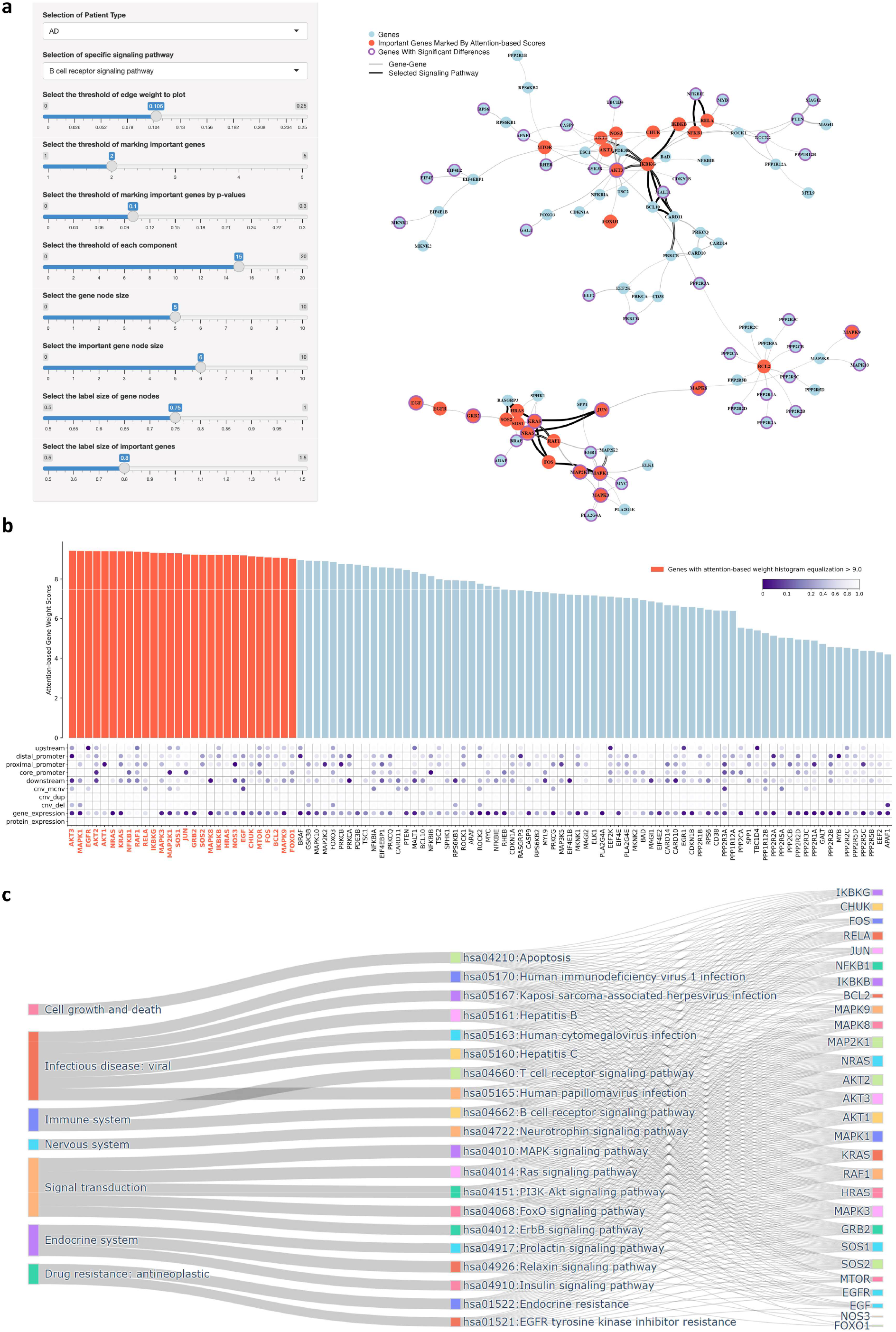
Biomarkers and pathways associated with Alzheimer’s disease. **a**. Visualization tool ***NetFlowVis*** for core signaling network interactions of AD. The important nodes are set as red filtered by attention-based node weight threshold of 2.0 and nodes with significant (p values <0.1) differences between AD and non-AD samples are circled with purples **b**. Barplot of the attention-based importance score measured by ***M3NetFlow***. The gene names and bars marked with red are important genes selected by setting attention-based node weight as 2.0, and the borders of the bar are set as purple if significant (p values <0.1) differences existed between AD and non-AD samples. **c**. Sankey plot of enriched signaling pathways of the identified AD associated genes selected by attention-based node weight (threshold as 2.0).

Among them, several signaling pathways play critical roles in Alzheimer’s disease (AD), particularly through mechanisms involving inflammation, immune responses, and cellular growth and death. For instances, the B cell receptor (BCR) and T cell receptor (TCR) signaling pathways are vital for B cell and T cell activation, which plays a crucial role in immune surveillance and inflammation. Shared molecular components between the BCR and TCR pathways, including MAP2K1, JUN, CHUK, RELA, NFKB1, IKBKB, and AKT genes, suggest significant cross-talk that contributes to neuroinflammatory processes in AD. Prolonged activation of these immune pathways may exacerbate chronic inflammation and contribute to AD pathology by sustaining harmful neuroinflammatory responses^45^. The NF-κB signaling axis, a critical component in both BCR and TCR pathways, is particularly relevant in AD due to its role in regulating inflammatory cytokine production. Chronic activation of NF-κB has been linked to increased amyloid-beta (Aβ) deposition and neurofibrillary tangle formation, both of which are hallmark features of AD pathology^46,47^. Additionally, activation of NF-κB, MAPK, and AKT signaling cascades can induce neuronal apoptosis, leading to cognitive decline associated with the disease.

The MAPK, RAS, PI3K-Akt, and FoxO signaling pathways also play critical roles in modulating immune responses and neuronal survival. The p38 MAPK pathway, for example, drives neuroinflammation by activating microglia and promoting the release of pro-inflammatory cytokines, which can result in neuronal death^48^. Dysregulation of the RAS-RAF-MEK-ERK signaling cascade, another crucial pathway in AD, affects neuronal survival and apoptosis. Overactivation of RAS signaling increases oxidative stress, contributing to Aβ accumulation and tau pathology, both of which are central to neurodegeneration in AD^49^. While Aβ accumulation is an early event in AD, tau pathology is more closely associated with cognitive decline, and together, these processes are considered the primary drivers of neuronal death in AD^50,51^.

The Akt signaling pathway, which promotes cell survival by inhibiting apoptosis through downstream effectors such as BCL2, is also impaired in AD. Reduced Akt activity has been linked to increased neuronal apoptosis and the accumulation of Aβ and hyperphosphorylated tau. Notably, the PI3K-Akt pathway is intimately connected to insulin signaling, which is disrupted in AD and is characterized by reduced Akt signaling, impaired glucose metabolism, and an increased vulnerability to neurodegeneration^52^. Furthermore, dysregulation of the FoxO transcription factors due to impaired PI3K-Akt signaling leads to enhanced expression of pro-apoptotic genes such as BIM and PUMA, contributing to synaptic and neuronal loss^53^.

Neurotrophin signaling, which regulates axonal growth and regeneration via MAPK and PI3K-Akt pathways, is another pathway that becomes disrupted in AD. This disruption impairs axonal repair mechanisms and contributes to synaptic loss and neuronal degeneration^54^. Additionally, endocrine-related pathways, including insulin and relaxin signaling, play crucial roles in maintaining cellular homeostasis and survival. Dysregulation of these pathways in AD contributes to neuronal apoptosis, neuroinflammation, and impaired protein clearance, further exacerbating disease pathology^52,55,56^.

## 4. Discussion and conclusion

Along with the advancement of next-generation sequencing (NGS) technology, multi-omic data of diseases are being generated to characterize the dysfunctional molecular targets and signaling pathways of complex diseases, like cancer and AD, which are valuable and essential for the development of personalized medicine or precision medicine prediction. However, it remains a challenging task to identify key molecular targets and core signaling pathways from a large signaling network with thousands of signaling targets and extensive signaling interactions. Herein, we present a novel graph AI model, named M3NetFlow, for generic multi-omic data integrative analysis. In this model, multi-omic data are used as the numerical features of individual proteins, and the protein-protein interactions form a large-scale signaling graph. Unlike existing graph models, we divided the large-scale network into signaling modules and applied a multi-hop strategy to better infer the essential proteins and their interactions. The experimental evaluation on two real multi-omic data analysis applications showed that the proposed model outperformed existing graph models in accuracy and is able to identify essential targets and core signaling pathways, demonstrating its interpretability for Alzheimer’s disease or drug combination synergy. The proposed model can be applied to 1) anchor-target guided or 2) generic target and pathway inference, as indicated by the two multi-omic data analysis applications that represent widely conducted tasks in omic data analysis. In the first scenario, a set of targets of interest, such as drug targets or known molecular targets, are used as the anchor nodes. Then, the signaling flows/information from neighboring graph nodes propagate to the anchor nodes. The embeddings of the anchor nodes are used as input for decoders to predict sample classes or drug combination responses. In the second scenario, the embeddings of graph nodes are pooled together as input for decoders to predict sample classes. Based on the attention scores of graph nodes, the essential targets and signaling pathways are further identified. Therefore, the proposed model can be applied for generic, integrative, and interpretable multi-omic data analysis tasks, and the code is publicly accessible.

It is still an exploratory study for multi-omic data analysis. There are some limitations that need further investigation. For example, more signaling pathways and larger protein-protein interactions should be evaluated. Moreover, dividing large signaling graphs into subnetworks or network modules can be achieved by using biologically meaningful annotations, such as gene ontology (GO) terms. Additionally, more multi-omic datasets are being generated. Combining multi-omic data from different diseases can provide a larger sample size than individual disease datasets, which could improve the training or pre-training of graph AI models and help identify pan-disease or disease-specific targets. It is also interesting to expand graph models from tissue-level multi-omic data to single-cell multi-omic data, which can be more challenging due to the large number of single-cell samples. Therefore, novel and improved graph AI models are needed to integrate and interpret multi-omic datasets, identify and infer key molecular targets and signaling pathways of complex diseases, and guide the development of precision medicine.

## Appendix

### Section A. Data collection and preprocessing

#### A.1 Drug combination screening datasets

In this paper, we utilized the NCI ALMANAC and O’Neil^57^ drug screening datasets to train the deep learning models. The NCI ALMANAC dataset consists of combo scores for various permutations of 104 FDA-approved drugs, representing their impact on tumor growth in NCI60 human tumor cell lines. For evaluating the synergy score of two drugs on a specific tumor cell line, we utilized the average combo-score of two drugs at different doses, employing a 4-element tuple: <*D*_*A*_, *D*_*B*_, *C*_*C*_, *S*_*ABC*_>. On the other hand, the O’Neil dataset, obtained from the DrugComb^58^ platform, served as another drug combination screening dataset. We extracted the processed dataset from this platform, which provided average synergy Loewe scores. These scores assessed the synergy score of two drugs on a given tumor cell line, employing a 4-element tuple: <*D*_*A*_, *D*_*B*_, *C*_*C*_, *S*_*ABC*_>.

#### A.2 Multi-omic data of cancer cell lines

In this study, multi-omic data comprising RNA-seq, copy number variation, gene methylation, and gene mutation data for a total of 1,489 genes were incorporated into the model. These data were obtained from the Cell Model Passports^59^ and CCLE^60^ databases. Through the identification of overlapping cell lines from these databases, we identified 42 cell lines that were present in both the Cell Model Passports database and the NCI ALMANAC dataset, which are A498, A549/ATCC, ACHN, BT-549, CAKI-1, DU-145, EKVX, HCT-116, HCT-15, HOP-62, HOP-92, HS 578T, IGROV1, K-562, KM12, LOX IMVI, MCF7, MDA-MB-231/ATCC, MDA-MB-468, NCI-H23, NCI-H460, NCI-H522, OVCAR-3, OVCAR-4, OVCAR-8, PC-3, RPMI-8226, SF-268, SF-295, SF-539, SK-MEL-2, SK-MEL-28, SK-MEL-5, SK-OV-3, SNB-75, SR, SW-620, T-47D, U251, UACC-257, UACC-62, UO-31. Additionally, we found 24 cell lines that were present in both the Cell Model Passports database and the O’Neil dataset, which are A2058, A2780, A375, CAOV3, HCT116, HT144, LOVO, MDAMB436, NCI-H460, NCIH1650, NCIH2122, NCIH23, OV90, OVCAR3, RKO, RPMI7951, SK-OV-3, SKMEL30, SKMES1, SW-620, SW837, T-47D, UACC62, VCAP.

#### A.3 Multi-omic data and clinical phenotypes datasets from ROSMAP

Following the acquisition of the datasets, they were reformatted into 2-dimensional data frames, with columns dedicated to sample identifiers, such as IDs and names, and rows corresponding to probes, gene symbols, and gene IDs. To successfully integrate the multi-omic data with clinical information, it was essential to match identical samples across the various datasets. This required standardizing the row data—including probes, gene symbols, and gene IDs—into a unified gene-level format, either by aggregating gene-specific measurements or by resolving duplicates from gene synonyms. Genes were subsequently mapped to a reference genome to ensure precise annotation within the multi-omic datasets. Gene counts were then normalized across the datasets, with missing values imputed using zeros or negative ones as needed. After aligning all columns to standard sample IDs and rows to standardized gene IDs, and ensuring a consistent number of samples and genes, the data was prepared for integration into Graph Neural Network (GNN) models, where epigenomic, genomic, transcriptomic and proteomic data were employed as features for nodes.

#### A.4 KEGG Signaling Pathways

About 59,241 gene-gene interactions for over 8,000 genes across different signaling pathways were collected from KEGG^61^ database. And 48 signaling pathways in the KEGG dataset were selected as the ground truth of subgraphs in the gene-gene interactions, which were AGE-RAGE signaling pathway in diabetic complications, AMPK signaling pathway, Adipocytokine signaling pathway, Apelin signaling pathway, B cell receptor signaling pathway, C-type lectin receptor signaling pathway, Calcium signaling pathway, Chemokine signaling pathway, ErbB signaling pathway, Estrogen signaling pathway, Fc epsilon RI signaling pathway, FoxO signaling pathway, Glucagon signaling pathway, GnRH signaling pathway, HIF-1 signaling pathway, Hedgehog signaling pathway, Hippo signaling pathway, Hippo signaling pathway - multiple species, IL-17 signaling pathway, Insulin signaling pathway, JAK-STAT signaling pathway, MAPK signaling pathway, NF-kappa B signaling pathway, NOD-like receptor signaling pathway, Neurotrophin signaling pathway, Notch signaling pathway, Oxytocin signaling pathway, PI3K-Akt signaling pathway, PPAR signaling pathway, Phospholipase D signaling pathway, Prolactin signaling pathway, RIG-I-like receptor signaling pathway, Rap1 signaling pathway, Ras signaling pathway, Relaxin signaling pathway, Signaling pathways regulating pluripotency of stem cells, Sphingolipid signaling pathway, T cell receptor signaling pathway, TGF-beta signaling pathway, TNF signaling pathway, Thyroid hormone signaling pathway, Toll-like receptor signaling pathway, VEGF signaling pathway, Wnt signaling pathway, cAMP signaling pathway, cGMP-PKG signaling pathway, mTOR signaling pathway, p53 signaling pathway. After preprocessing the dataset, 17,259 gene-gene interactions for 1,489 genes were obtained. And 28,843 gene-gene interactions for 2,144 genes were obtained for ROSMAP AD dataset.

#### A.5 Drug-Target interactions derived from DrugBank

Drug-target information was extracted from the DrugBank^62^ database (version 5.1.5, released 2020-01-03). In total, 15,263 drug-target interactions were obtained for 5435 drugs/investigational agents and 2775 targets. Further, 17 drugs with known targets to the 1489 genes previously identified were selected for use in our model for NCI ALMANAC^63^, specifically: Celecoxib, Cladribine, Dasatinib, Docetaxel, Everolimus, Fulvestrant, Gefitinib, Lenalidomide, Megestrol acetate, Mitotane, Nilotinib, Paclitaxel, Romidepsin, Sirolimus, Thalidomide, Tretinoin, Vorinostat. And 10 drugs with known targets to the 1489 genes previously identified were selected for use in our model, specifically: Dasatinib, Erlotinib, Lapatinib, Sorafenib, Sunitinib, Vorinostat, Geldanamycin, Metformin, Paclitaxel, Vinblastine.

### Section B. Method Details

#### B.1 *K*-hop attention-based graph neural network

To incorporate the long-distance information, for each subgraph *K* -hop attention based mechanism was constructed with

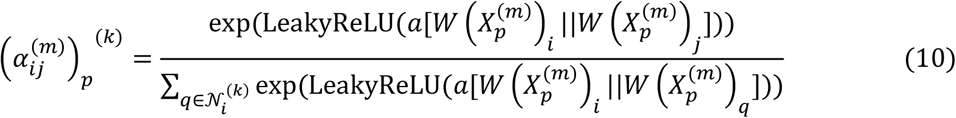

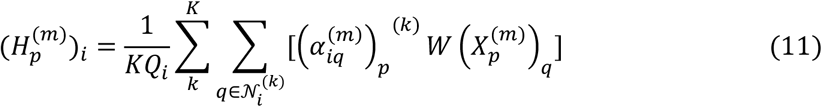

where 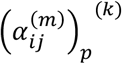 is the attention score mentioned in equation (1) and 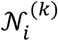 is the neighbor nodes calculated by the adjacency matrix in *k*-th hop *A*^(*k*)^ for the node *i* (See **Appendix B.3** for the algorithm of *k*-th hop adjacency matrix). The linear transformation vector *a* ∈ ℝ^2*d’*^was also defined. At the same time, the linear transformation for features of each node will be defined as *W* ∈ ℝ^*d*×*d*’^. And 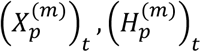 represent the feature and updated feature of node *t* (*t* =1, 2, …, *n*) . Aside from that, *Q*_*t*_ represents the number of signaling pathway the node *t* (*t* =1, 2, …, *n*) belongs to. And above formula shows the calculation for the 1-head attention. The number of head *h* will be modified in the model, and the node embeddings for each subgraph will take the average of embeddings in every head attention.

#### B.2 Weighted Bi-directional Message Propagation

The weight matrices are represented as 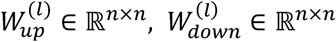 for each of the *L* global message propagation layers, (*l* = 1, 2, …, *L*). The weighted adjacency matrices are defined as follows:

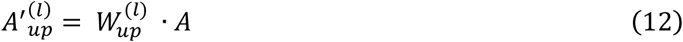

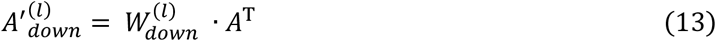

Here,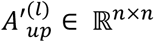 and 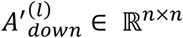 maintain the parameters for drug-gene relationships as in the original adjacency matrix. The mean aggregation of each node’s neighbors’ features from upstream to downstream from layer *l* ™ 1 to layer *l* is achieved using the following equation:

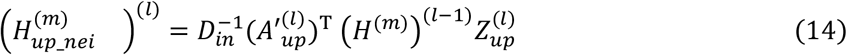

Similarly, the mean aggregation of each node’s neighbors’ features from downstream to upstream is computed as:

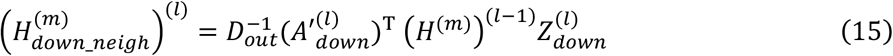

The biased transformation of nodes and their features is given by:

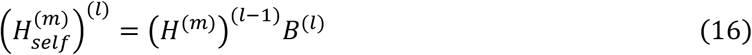

In these equations, 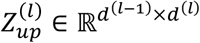 and 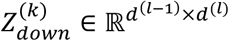 are the linear transformation matrices for each layer *l*, (*l* = 1, 2, …, *L*). For each layer, the model employs a normalization function 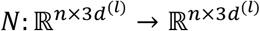 and a LeakyReLU activation function with parameter α to map the concatenated node features 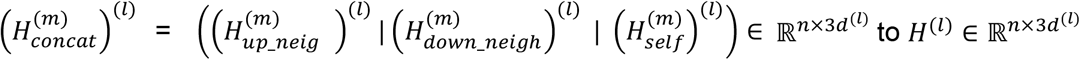 as follows

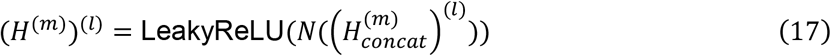

where the normalization function *N* performs L2 normalization on the demo matrix *V* ∈ ℝ^*p*×*q*^ along the row axis using the equation:

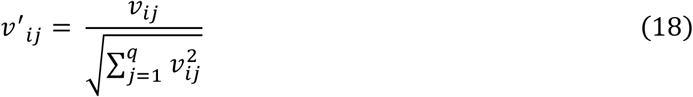

where *v*’_*ij*_ is an element of the new matrix *V*’ ∈ ℝ^*p*×*q*^.

#### B.3 *K*-hop adjacency matrix generation algorithm

##### Algorithm

***k***-th hop adjacency matrix

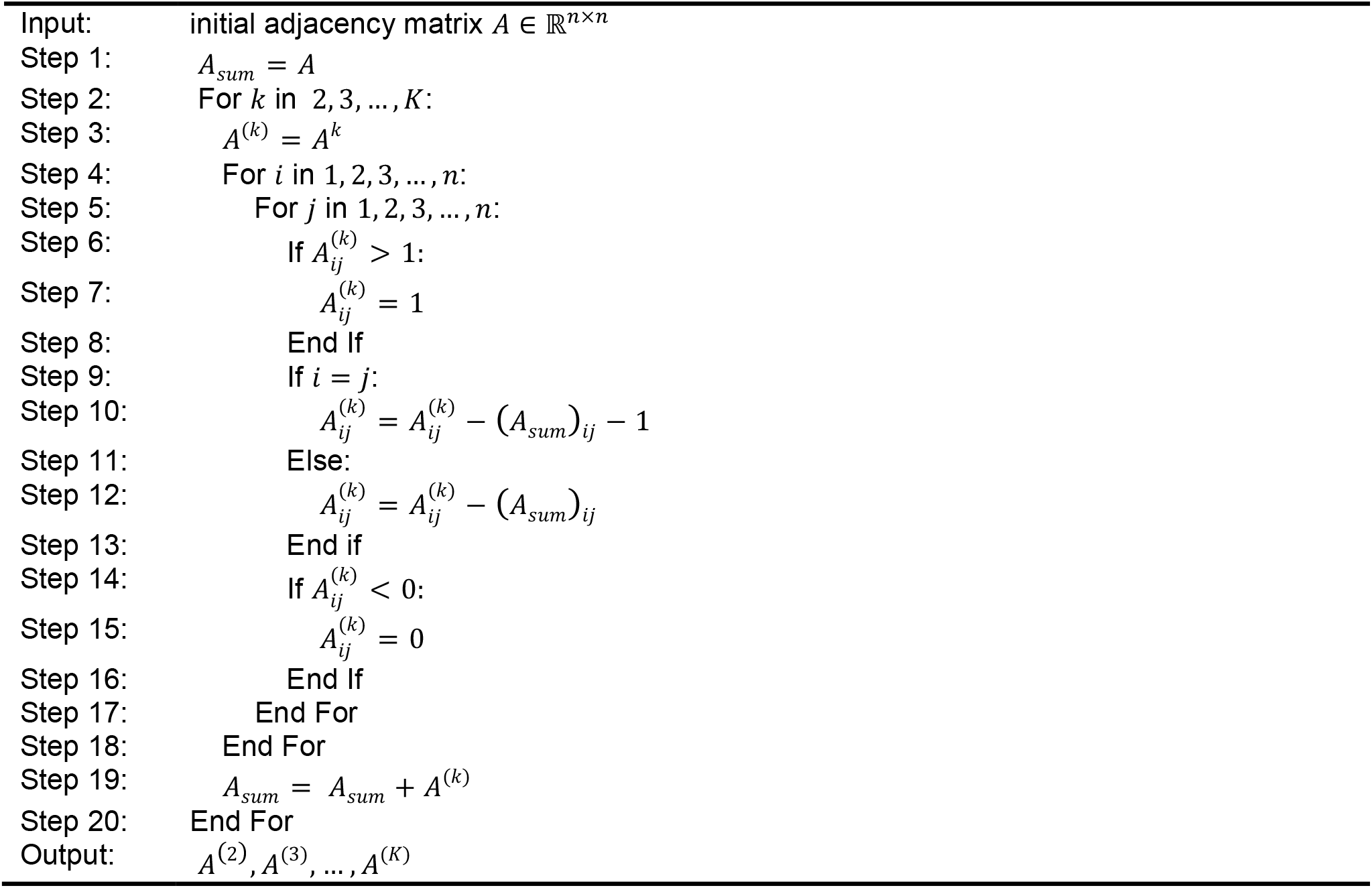

##### Section C. Downstream analysis of important targets for cancer cell lines

**Figure S1.**
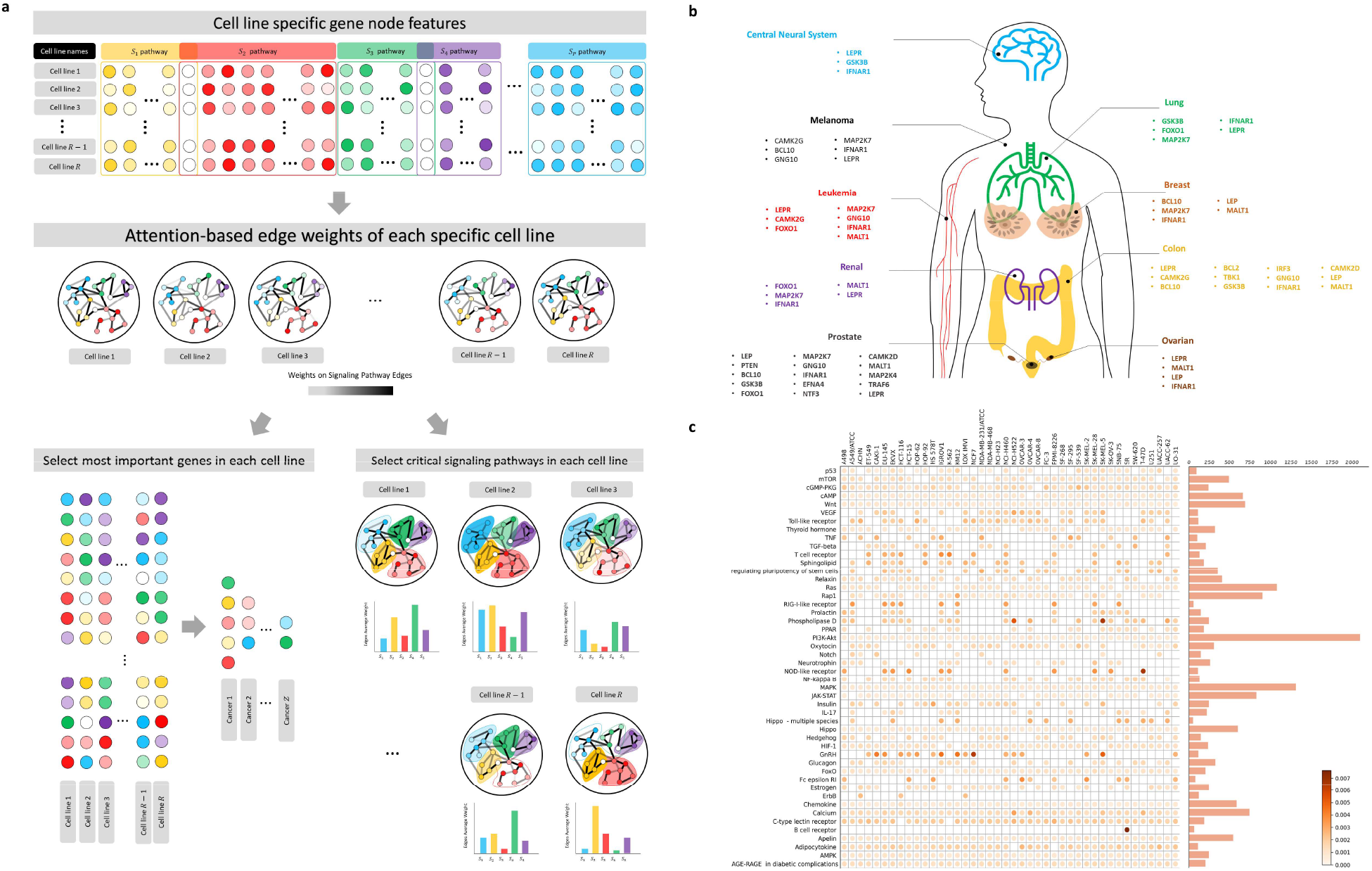
Downstream analysis of important genes and signaling pathways. a. Procedures of identifying the important genes in different cell lines and cancers, and identifying the important signaling pathways in different cell lines. b. Overlapped genes for the cell lines in each type of cancer. c. Heatmap of sum of edge weights in different signaling pathways in whole NCI ALMANAC dataset.

##### Section D. *NetFlowVis*: a tool to visualize core signaling pathways associated with synergistic drug combinations

To visualize the results and gain a better understanding of the underlying mechanism of drug effects, we generated a core network of signaling interactions by applying a threshold. To enhance the interactions, we utilized the RShiny package and developed a visualization tool called ***NetFlowVis*** (The website link has been provided in the code availability part). This tool allows users to control the edge threshold and set a minimum number of nodes in each network component. Users have the option to select the desired cell line for visualization and mark specific signaling pathways for detailed analysis. As an example, **Figure 3b** demonstrates the visualization of the cell line DU-145. Furthermore, by referring to the top 20 prior gene targets and their corresponding gene degree (node importance scores) for cell line DU-145, we observed that the majority of gene targets were included in the filtered signaling network interactions.

By utilizing the ***NetFlowVis*** visualization tool, users could take control of the size of the core signaling network. In the case of the DU-145 cell line, an edge threshold of 0.31 was selected, which corresponded to approximately the 98.5th percentile for edge weight among all edges. Additionally, a threshold of 23 was chosen for marking important nodes/genes in red, representing ∼99.9th percentile for node importance scores across all nodes. To gain a deeper understanding of the underlying mechanism, the interactions between drugs and genes were analyzed, with the highest and lowest drug-gene interactions being represented by the colors purple and green, respectively. The analysis revealed that the drug ***Paclitaxel*** targeted the important gene ***BCL2***, while the drug ***Celecoxib*** targeted less essential genes (see **Figure 3b**). This finding demonstrated that targeting the important genes selected by the ***M3NetFlow*** model led to higher drug scores, as depicted in **Figure 3c-f**. Moreover, by choosing a specific signaling pathway within the core signaling network, it was observed that the ***IL-17 signaling pathway*** was downstream of the critical gene ***TRAF6***. Interestingly, experimental validation has shown the involvement of the ***IL-17 signaling pathway*** in promoting migration and invasion of ***DU-145*** cell lines through the upregulation of ***MTA1*** expression in prostate cancer^64^.

##### Section E. Code Availability

*M3NetFlow* code: https://github.com/FuhaiLiAiLab/M3NetFlow

*NetFlowVis* app: https://m3netflow.shinyapps.io/NetFlowVis/

## Acknowledgment

This study was partially supported by NIA R56AG065352 (to Li), NIA 1R21AG078799-01A1 (to Li/Province), NINDS 1RM1NS132962-01 (to Dickson/Marco/Cooper/Li),Children’s Discovery Institute (CDI) M-II-2019-802, and NLM 1R01LM013902-01A1 to Li.

## Author contributions

FL conceived this study. FL, HZ designed the model and prepared the manuscript. HZ implemented the model and analyzed the data and results. FL, HZ, PG, LD, WK, DD, RF, YC, PP discussed the method and results.

